# Adaptive iron utilization compensates for the lack of an inducible uptake system in *Naegleria fowleri* and represents a potential target for therapeutic intervention

**DOI:** 10.1101/763011

**Authors:** Dominik Arbon, Kateřina Ženíšková, Jan Mach, Maria Grechnikova, Ronald Malych, Pavel Talacko, Robert Sutak

**Author notes:** Corresponding author (RS).

## Abstract

Warm fresh water is a natural habitat for many single-celled organisms, including protozoan parasites such as the infamous brain-eating amoeba, *Naegleria fowleri,* which can become pathogenic for mammals, including humans. The condition caused by *N. fowleri* is known as primary amoebic meningoencephalitis, which is a generic and usually fatal infection of the brain with rapid onset. One of the important factors influencing a wide spectrum of pathogens, including *N. fowleri*, is the bioavailability of iron in the environment. The strategy of withholding iron evolved in mammalian hosts, and the different strategies of pathogens to obtain it are an important part of host-parasite interactions.

In the present study, we employ different biochemical and analytical methods to explore the effect of decreased iron availability on the cellular processes of *N. fowleri*. We show that under iron starvation, nonessential, iron-dependent, mostly cytosolic pathways in *N. fowleri* are downregulated, while the metal is utilized in the mitochondria to maintain vital respiratory processes. Surprisingly, *N. fowleri* fails to respond to acute shortages of iron by the induction of a reductive iron uptake system that seems to be the main iron-obtaining strategy of the parasite. Our work aims to demonstrate the importance of mitochondrial iron in the biology of *N. fowleri* and to explore the plausibility of exploiting it as a potential target for therapeutic interference.

**Author Summary:** *Naegleria fowleri* is undoubtedly one of the deadliest parasites of humans, hence the name “brain-eating amoeba”. Being dangerous, but rare, it may be regarded as a highly understudied pathogen of humans. Unfortunately, the symptoms of primary amoebic meningoencephalitis may be confused with much more common bacterial meningoencephalitis; therefore, it is quite probable that many infections caused by *N. fowleri* have been misdiagnosed as bacterial infections without further inquiry. In many cases, fast diagnosis is vital for commencing the correct therapy, and even then, complete success of the treatment is very rare. Our laboratory focuses on the uptake and intracellular metabolism of metals in unicellular eukaryotes, so we decided to explore the biology of *N. fowleri* from this aspect. Changes in the proteome, as a direct effect of iron-deficient conditions, were described, and these data were used to further explore the ways in which *N. fowleri* responds to these conditions on a cellular level and how its biology changes. Based on these findings, we propose that the struggle of *N. fowleri* to obtain iron from its host could be exploited for therapeutic interference purposes in primary amoebic meningoencephalitis patients.

## Introduction

There are several amoebae that, in certain conditions, are able to parasitize suitable hosts. One particular member of this group, *Naegleria fowleri*, is widely known as the “brain-eating amoeba”. This rather vivid nickname is attributed to the condition known as primary amoebic meningoencephalitis (PAM), which is caused by *N. fowleri*. It is found in warm fresh waters across most continents and can alter between three distinguishable forms: a durable cyst, trophozoite amoeba and mobile flagellar form [1]. Its presence is dangerous for people partaking in recreational activities involving water bodies [2]. The disease is acquired when *N. fowleri* trophozoites forcefully enter through the nasal cavity and intrude the olfactory neuroepithelium [3]. The disease has a rapid onset, and the nonspecific symptoms resemble those of bacterial meningoencephalitis – headache, fever, vomiting – with rapid progress, causing seizures, coma, and death [4,5]. Since PAM occurs commonly in healthy individuals, *N. fowleri* is not regarded as an opportunistic parasite but as a pathogen [3]. Several other nonpathogenic species of *Naegleria* exist, and *Naegleria gruberi* and perhaps *Naegleria lovaniensis* receive particular interest because these strains are generally accepted as suitable laboratory systems [6].

The fatality rate for PAM is reported to be above 97% [7]. Its treatment is difficult not only because of the interchangeable symptoms and rapid onset but also because there is no specific medication for the disease. For successful treatment, it is crucial to make a correct diagnosis in time, followed by immediate therapy. The currently accepted treatment includes administering several medications simultaneously, such as amphotericin B, fluconazole, rifampin, azithromycin or miltefosine, in combination with methods that decrease brain swelling [4,8].

Iron is an essential constituent of many biochemical processes, including redox reactions, the detoxification of oxygen, cell respiration or various enzymes. This is mainly due to its ability to alter between different oxidation states, therefore partaking in redox reactions, often in the form of iron-sulfur clusters [9]. Iron is essential for virtually all known forms of life, and its availability is shown to be the limiting factor of growth in certain locations [10]. Iron acquisition becomes even more challenging for pathogens that inhabit another living organism, as demonstrated in the case of *Plasmodium*, where iron-deficient human and mouse models manifested unfavorable conditions for parasite development [11,12]. The importance of iron acquisition mechanisms is known for other parasites as well [13]. Similarly, it is suggested that iron-deficient conditions have an adverse effect on the viral life cycle [14], and several bacteria are known to produce iron-chelating compounds, siderophores, to take up iron from their surroundings. As a defense mechanism, mammals minimize the presence of free iron in their body using several proteins, such as lactoferrin, ferritin or transferrin, which bind the metal, decreasing its bioavailability [15]. Parasitic organisms are known to have adapted various means of acquiring iron from their environments, ranging from opportunistic xenosiderophore uptake to specific receptors for mammalian iron-containing proteins [13]. This two-sided competition between pathogens and their hosts indicates the importance of iron metabolism in disease and underlines the importance of further research on this topic to search for new methods of therapeutic intervention.

This work demonstrates how *N. fowleri* struggles to successfully adapt to iron-deficient conditions. No increase in iron uptake from its surroundings, either directly or through phagocytosis, was detected, and no alternative metabolic pathway was shown to adjust for the condition. Proteomics and metabolomics approaches showed that the response of *N. fowleri* to low iron levels is to maintain iron trafficking to mitochondria, while nonvital cytosolic iron-requiring pathways decline. These findings reveal a possible exploitable weakness in the survival strategy of the amoeba within its host. Although the host brain is relatively rich in iron content [16], not all of the iron is readily available for the parasite, as we show in the case of iron bound to transferrin, the main human extracellular iron-binding protein. Thus, disrupting the process of iron uptake by *N. fowleri* in the human brain, such as using artificial chelators, could represent a safe complementary antiparasitic strategy against this deadly pathogen.

## Results

### Iron chelators have the potential to significantly hinder *N. fowleri* growth

Iron plays a vital role in many biochemical processes; therefore, it is rational to expect that decreasing the bioavailability of iron in the surrounding environment hinders cell growth. To determine the effect of iron on the cell propagation of *N. fowleri*, three different iron chelators were added to *N. fowleri* growth medium: bathophenanthroline disulfonic acid (BPS), 2,2′-dipyridyl (DIP) and deferoxamine (DFO). The compounds inhibited the growth of the cultures to different extents. The IC_50_ values were calculated for each compound after cultivation for 48 hours and are summarized in Table 1. The most potent effect in hindering the culture growth was observed with the siderophore DFO. Both BPS and DIP exhibited notably higher IC_50_ values. For all further experiments presented in this work, unless stated otherwise, iron deprivation was achieved using 25 μM of the common extracellular chelator BPS, a condition when cell growth was significantly affected, but microscopy observation showed no encystation or flagellated form, and the cells did not lose their ability to multiply or attach to the surface. This is an important prerequisite, particularly for proteomic and transcriptomic analysis, where too complex changes are accompanied by unfavorable cell processes that could bias the data. To achieve iron-rich conditions, the cultivation medium was supplemented with 25 μM ferric nitrilotriacetate (Fe-NTA).

**Table 1.**
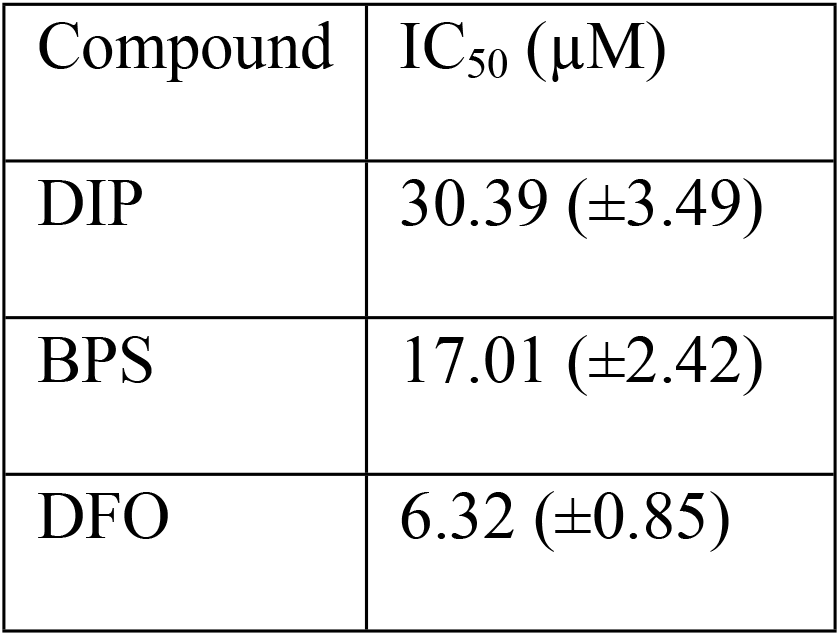
Iron chelator IC_50_ values for *N. fowleri* cultures after 48 hours.

| Compound | IC<sub>50</sub> (µM) |
| --- | --- |
| DIP | 30.39 (±3.49) |
| BPS | 17.01 (±2.42) |
| DFO | 6.32 (±0.85) |

### Iron uptake is limited by ferric reductase activity

Iron is generally available in the environment in two distinct oxidation states. Due to the different solubilities of the two forms, many organisms use ferric reductase to reduce ferric to ferrous iron, which is more soluble and therefore easier to acquire. To explore the mechanism by which *N. fowleri* acquires iron from its surroundings, the cell cultures were supplemented with different sources of iron: ^55^Fe-transferrin, ^55^Fe(III)-citrate and ^55^Fe(II)-ascorbate. The results presented in Fig 1A show that ferrous iron is taken up and incorporated into intracellular protein complexes more rapidly than its trivalent counterpart, suggesting the involvement of ferric reductase in the effective uptake and utilization of iron. The insignificant uptake of transferrin shown in S1 Fig (A) shows that this protein is not a viable source of iron for *N. fowleri*, corresponding to the fact that it is not found in the usual habitat of this organism. Surprisingly, there was no observable difference in iron uptake between cells precultivated in iron-rich or iron-deficient media, showing that the organism fails to stimulate its iron uptake machinery.

**Fig 1.**
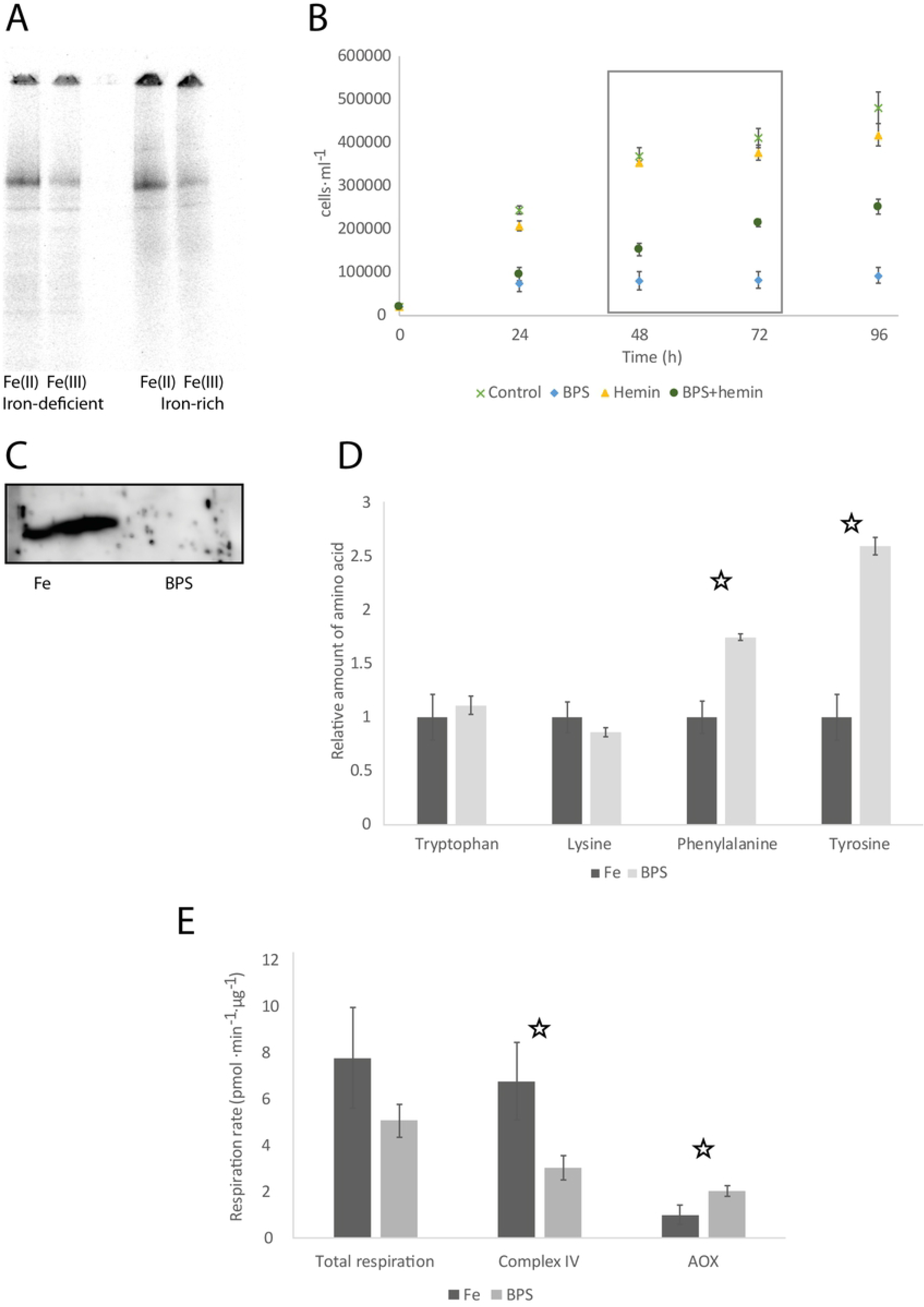
Effect of iron deficiency on *N. fowleri*. **(A) Ferrous and ferric iron uptake by *N. fowleri* precultivated in iron-rich and iron-deficient conditions.** Autoradiography of blue native electrophoresis gels of whole cell extracts from *N. fowleri* previously cultivated for 72 hours in iron-deficient conditions (25 μM BPS) or iron-rich conditions (25 μM Fe-NTA) and further incubated with ^55^Fe(II) (ferrous ascorbate) and ^55^Fe(III) (ferric citrate). Equal protein concentrations were loaded. **(B) Growth restoration of *N. fowleri* culture by hemin in iron-deficient conditions.** Cells grown in regular growth medium (control) and with 50 μM hemin (hemin) exhibited similar propagation, while cells in iron-deficient conditions achieved with 50 μM chelator bathophenanthroline disulfonic acid (BPS) were stagnate in growth in the first 24 hours. The addition of 50 μM hemin to the iron-deficient conditions (BPS+hemin) partially restored culture growth. Boxed area signifies time, where growth restoration was calculated from. Data are presented as means ± SD from three independent experiments. **(C) Downregulation of *N. fowleri* hemerythrin in iron-deficient conditions.** Western blot analysis of cells cultivated in iron-rich (Fe) and iron-deficient (BPS) conditions using an antibody against *Naegleria* hemerythrin. The amount of protein loaded was equalized. **(D) Cellular content of selected amino acids in *N. fowleri* cells cultivated in iron-rich and iron-deficient conditions.** Relative amounts of phenylalanine and tyrosine were significantly increased in the iron-deficient condition, while tryptophan and lysine remained unchanged. T-test p-values <0.01 are marked with a star. The total protein concentration was equal in all samples, and values are given relative to the iron-rich conditions for each amino acid. Fe, cells cultivated in iron-rich conditions; BPS, cells cultivated in iron-deficient conditions. Data are presented as means ± SD from three independent experiments. **(E) Respiration of *N. fowleri* grown in iron-rich and iron-deficient conditions.** Using selective inhibitors of complex IV and alternative oxidase, the contribution of alternative oxidase and complex IV activity to the total respiration was assessed. AOX, alternative oxidase; Fe, cells cultivated in iron-rich conditions; BPS, cells cultivated in iron-deficient conditions. The rates of respiration are shown relative to the values of cells cultivated in iron-rich conditions. The T-test p-values <0.01 are marked with a star. Data are presented as means ± SD from five independent experiments.

To confirm that *N. fowleri* employs a reductive iron uptake mechanism, cell cultures were presented with a source of a ferric iron radionuclide in the presence and absence of the ferrous iron chelator BPS. The results shown in S1 Fig (B) demonstrate that *N. fowleri* readily incorporates ferric iron into its protein complexes, while BPS chelates initially reduced ferrous iron and prevents it from being further utilized, confirming that ferric iron uptake requires the reduction step.

### Ferric reductase activity is not affected by iron availability

We have shown that neither ferrous nor ferric iron uptake is induced by iron starvation (Fig 1A). To further investigate this observation, the activity of ferric reductase was assessed in cells preincubated in iron-rich and iron-deficient conditions. Measuring ferric reductase activity showed that the conversion of ferric to ferrous iron was not significantly changed in iron-deprived cells. This further confirms that *N. fowleri* cannot efficiently adjust the rate of iron acquisition from its surroundings in dependence on iron availability.

### Hemin partially restores *N. fowleri* growth in the iron-deficient medium

As shown in Fig 1B, cells cultivated in regular growth medium have a similar propagation pattern in comparison to a culture grown in medium supplemented with 50 μM hemin; therefore, at a given concentration, hemin is not toxic for *N. fowleri*, nor does it significantly support its growth under standard cultivation conditions. However, while 50 μM BPS arrested the culture growth at 31% (±12%) during the first 24 hours, the addition of 50 μM hemin partially restored culture growth to 42% (±10%) in 48 hours and up to 52% (±7%) by 72 hours. The T-test P-values for cultures in BPS and BPS supplemented with hemin are <0.01 for 48 and 72 hours of growth.

### The proteomic analysis illuminates the iron-starvation response of *N. fowleri*

Since iron has an irrefutable role as a cofactor for various enzymes and its metabolism is dependent on a great number of proteins, we aimed to examine the overall effect of iron availability on the *N. fowleri* proteome. Therefore, we compared the whole-cell proteomes of cells grown in iron-rich and iron-deficient conditions, and we have additionally analyzed membrane-enriched fractions of cells to obtain a higher coverage of the membrane proteins. The aim was to reveal the metabolic remodeling accompanying iron starvation and to identify proteins responsible for iron homeostasis, such as membrane transporters, signaling and storage proteins or proteins involved in iron metabolism, for example, the formation of iron-sulfur clusters.

S1 Table lists the proteins that were significantly upregulated or downregulated in iron-deficient conditions based on the analysis of the *N. fowleri* whole-cell proteome, and S2 Table contains the regulated proteins in the membrane-enriched fraction. The proteins most relevant for this work are summarized in Table 2. The list of downregulated proteins based on whole-cell proteomics under iron-deficient conditions contained 20% of predicted iron-containing proteins, most of which were nonheme enzymes such as desaturases and oxygenases, or hydrogenase (NF0008540) and its maturases HydE (NF0081220) and HydG (NF0081230). Importantly, most of the downregulated iron-containing proteins are typically located outside mitochondria.

**Table 2.** Effect of iron deprivation on the abundance of selected *N. fowleri* proteins.

| Gene ID | Protein | Whole cell (membrane fraction)<br>proteomics fold-change in iron-deficiency |
| --- | --- | --- |
| NF0008540 | Hydrogenase | -3.7 |
| NF0081220 | HydE | -2.8 |
| NF0081230 | HydG | -2.4 |
| NF0127030 | Hemerythrin | -7045.1 |
| NF0119700 | Hemerythrin | -11.4 |
| NF0117840 | Protoglobin | Infinite downregulation |
| NF0106930 | Phenylalanine hydroxylase | -5.7 |
| NF0084900 | 4-Hydroxyphenylpyruvate dioxygenase | -1.5 |
| NF0073220 | Homogentisate 1,2-dioxygenase | -1.8 |
| NF0001440 | Cysteine desulfurase | 3.7 |
| NF0060430 | IscU * | 3.0 |
| NF0001420 | Mitochondrial phosphate carrier protein * | 8.0 (8.4) |
| NF0079420 | Mitoferrin * | 1.8 (1.7) |
| NF0071710 | Deferrochelataase | 2.3 |
| NF0014630 | Mitochondrial carnitine/acylcarnitine transferase * | Not found (1.5) |
| NF0020800 | Fe superoxide dismutase | 1.8 |
| NF0044680 | Iron III Superoxide Dismutase | 1.1 |
| NF0030360 | Cu-Zn superoxide dismutase | 5.8 |
| NF0121200 | Catalase | -1.3 |
| NF0015750 | Manganese superoxide dismutase | 1.6 |
| NF0048730 | Glutathione peroxidase | 2.0 |
| NF0036800 | Iron-containing cholesterol desaturase daf-36-like | -2.0 (-3.4) |
| NF0001470 | Tyrosine aminotransferase | -2.0 |
| NF0004720 | Alternative oxidase | 1.5 |

Proteins were manually annotated using HHPRED or by sequence alignment with homologous proteins from other organisms (denoted with a star). The fold change based on proteomics analysis with the whole cells and membrane (for selected proteins; in brackets) fractions is given for cells cultivated in iron-deficient conditions. Significantly regulated proteins (with a fold change above 2.3) are highlighted in green.

The dramatic downregulation of the protein hemerythrin was confirmed by Western blotting using an antibody against *N. gruberi* hemerythrin. The result shown in Fig 1C shows that the corresponding band was diminished under iron-deficient conditions, which corresponds to the proteomics data. Ponceau S staining to confirm a uniform amount of loaded proteins is shown in S1 Fig (C).

In the phenylalanine catabolism pathway, all three iron-dependent enzymes, phenylalanine hydroxylase (NF0106930), p-hydroxyphenylpyruvate dioxygenase (NF0084900) and homogentisate 1,2-dioxygenase (NF0073220), were downregulated in iron-deprived cells, even though the decrease in the expression of the last two enzymes was below our set threshold.

The proteins upregulated under iron-deficient conditions included two essential mitochondrial components of iron-sulfur cluster assembly machinery, namely, cysteine desulfurase (NF0001440) and iron-sulfur cluster assembly enzyme IscU (NF0060430); two mitochondrial transporters, phosphate carrier (NF0001420) and iron-transporting mitoferrin (NF0079420); and a potential homologue of deferrochelatase (NF0071710). The increase in the expression of mitoferrin was below our set threshold, but the results from the comparative analysis of the membrane-enriched fraction of *N. fowleri* grown under different iron conditions (S2 Table and Table 2) confirmed the upregulation of this transporter as it did for the mitochondrial phosphate carrier. Additionally, in the membrane-enriched proteomics analysis, a carnitine/acylcarnitine transferase (NF0014630) was identified and found to be slightly, although not significantly, upregulated.

Iron metabolism is interconnected with the production and detoxification of reactive oxygen species. Among the antioxidant defense proteins, one family of superoxide dismutases contains iron as a cofactor. Our proteomic analysis showed that while iron-dependent SODs (NF0020800 and NF0044680) were not significantly changed, Cu-Zn-dependent SOD (NF0030360) was upregulated in iron-deficient conditions, suggesting a compensatory mechanism for the mismetallated iron-dependent enzyme. Other radical oxygen species detoxification enzymes, such as catalase (NF0121200), manganese SOD (NF0015750), or glutathione peroxidase (NF0048730), were not significantly regulated in iron-deficient conditions.

### Transcriptomics does not reflect the proteomic changes of *N. fowleri*

To uncover a broader spectrum of the affected proteins that we were unable to detect in proteomic analysis, we performed transcriptomic analysis of *N. fowleri* grown under the same conditions of iron availability (S3 Table). To our surprise, the obtained data did not correspond to the proteomics results. The only relevant genes regulated in the same way as the proteins observed by the proteomics analysis were hemerythrin (NF0127030, NF0119700), protoglobin (NF0117840) and iron-containing cholesterol desaturase daf-36-like (NF0036800). This result could indicate that the *N. fowleri* response to iron starvation is mostly posttranslational.

### The amino acid assay shows downregulation of the phenylalanine degradation pathway as a result of decreased iron availability

Considering the iron-dependent changes in the abundance of proteins participating in the phenylalanine degradation pathway, the next aim of this work was to analyze this effect by directly detecting metabolites, namely, quantifying the levels of corresponding amino acids in the cells by metabolomics. Cells grown in iron-rich and iron-deficient conditions were lysed, the protein concentrations were equalized, and the resulting materials were analyzed for amino acid content by liquid chromatography coupled with mass spectrometry.

As a control, relative amounts of tryptophan and lysine, the levels of which were not expected to change, were determined in the same samples. Relative amounts of both phenylalanine and tyrosine were significantly increased in iron-deficient conditions, as predicted by the proteomic data. The T-test values for tryptophan and lysine were 0.472 and 0.236, respectively; therefore, these values were not significant. However, the values for phenylalanine and tyrosine were 0.007 and less than 0.001, respectively, confirming the significant difference between the iron-deficient and iron-rich conditions. Data are shown in Fig 1D.

### Intracellular lactate production is decreased in iron-deficient cells

*N. fowleri* cells possess a lactate dehydrogenase and are therefore potentially able to produce lactate from pyruvate while replenishing the level of the cofactor NAD^+^. To analyze the effect of iron starvation on this metabolic pathway, lactate production was analyzed by gas chromatography coupled with mass spectrometry detection. The level of intracellular lactate in iron-deficient cells decreased by 32% (±13%, T-test p-value <0.05).

### Alternative oxidase activity shifts in iron-deficient conditions to preserve the total respiration rate

Alternative oxidase (AOX), present in *N. fowleri*, is a part of the mitochondrial respiratory chain. Its function lies in accepting electrons from ubiquinol to reduce the final electron acceptor, oxygen. Therefore, function of AOX is similar to complex IV; however, branching electron flow towards AOX bypasses some of the proton pumping complexes, decreasing the effectivity of the respiration chain. Using selective inhibitors of respiration complex IV and AOX enables the study of their participation in respiration. Here, the effect of decreased iron availability on the respiratory chain was depicted. The results shown in Fig 1E demonstrate that the total respiration of cells grown under iron-deficient conditions was decreased, although not significantly (T-test P-value >0.01), and this decrease was based on the lower activity of complex IV (T-test P-value <0.01). At the same time, in the iron-deficient cells, the activity of AOX significantly increased (T-test P-value <0.01).

### Phagocytosis reflects *N. fowleri* viability in iron-rich and iron-deficient conditions

Trophozoites of the amoeba feed on bacteria in their natural environment, presenting a potential source of iron. To investigate the overall cell viability and to test whether *N. fowleri* cells induce phagocytosis as an effort to stimulate the acquisition of a potential iron source, the ability of iron-starved cells to phagocytose bacteria was investigated. Quantification of phagocytosis was determined by flow cytometry using *Escherichia coli* that present increased fluorescence within the acidic endocytic compartments. Values were calculated as a percentage of amoeba containing phagocytosed bacteria from the total population. In iron-deficient conditions, the rate of phagocytosis was lower by 25% (±12%, T-test p-value <0.001) in comparison to iron-rich conditions, showing that iron availability plays a role in this process. A typical cell phagocytizing fluorescent bacteria is shown in S2 Fig.

## Discussion

Due to the irreplaceable role of iron in cellular processes, it is unsurprising that iron chelators have the ability to hinder the proliferation of *N. fowleri* [17]. The different chelators used in this work have distinct properties that influence their impact on cells. DIP is a membrane-permeable compound with affinity to ferrous iron [18] and a ratio of three chelator molecules binding one molecule of metal [19]. BPS is a membrane-impermeable chelator that binds ferrous iron with a ratio of three chelator molecules to one metal ion [20]. Finally, DFO is a membrane-impermeable, ferric iron-binding siderophore [21] with a binding ratio of one iron per molecule [22]. Quite surprisingly, the only tested membrane-permeable iron chelator, DIP, had the highest IC_50_ value (30.39 μM; SD = 3.49). In comparison with BPS, with which it shares an affinity to ferrous iron and the same denticity, DIP was almost half as effective in inhibiting culture growth. BPS and DFO differed not only by the oxidation state of bound iron but also by the ratio of molecules bound to the chelated iron. Their IC_50_ values were 17.01 μM (SD = 2.42) and 6.32 μM (SD = 0.85), respectively, showing an apparent tendency of the approximately three-fold amount of BPS required to have the same effect as DFO, probably because of the binding ratio. Therefore, surprisingly, it appears that from the selected chelators, the membrane-impermeable chelators are more effective against *N. fowleri.* It is important to consider that the growth medium used in this work was adjusted to the amoeba requirements, while in its host, the parasite is expected to meet much harsher conditions. Thus, the anticipated *in vivo* antiparasitic effect of suitable chelators may be more pronounced.

To maintain an optimal level of cellular iron in a hostile environment, such as host tissues, pathogens possess selective and effective mechanisms for iron uptake. These strategies of obtaining iron include the utilization of various sources from the host, including transferrin, lactoferrin or heme, and some parasites even exploit bacterial siderophores as sources of iron [13]. Transferrin, an abundant human blood protein that transports iron to various tissues, including the brain [23], serves as a viable source of iron for different parasites [13]. It was demonstrated that *N. fowleri* possesses proteases able to degrade human holotransferrin, although the work did not identify the intracellular fate of the iron [24]. Our work shows that *N. fowleri* does not have the means of efficiently utilizing iron from this host protein, perhaps because it is a facultative pathogen with no advantage of such an iron uptake mechanism in its natural environment. We further demonstrated the preference of *N. fowleri* for ferrous iron compared to ferric iron, the inhibitory effect of ferrous iron chelator on ferric iron uptake and the presence of extracellular ferric reductase activity. Based on these observations, we argue that the main strategy of iron acquisition by this parasite is the reductive two-step iron uptake mechanism, as described for *Saccharomyces cerevisiae* [13].

*S. cerevisiae* possesses the ability to upregulate the expression of ferric reductase, which is responsible for the first step of the reductive iron uptake mechanism, up to 55 times in iron-deficient conditions [25], therefore increasing the rate of iron uptake. Our work shows that in *N. fowleri*, the activity of ferric reductase is not induced by iron starvation nor is the ferric and ferrous iron uptake and further incorporation of iron into cellular proteins. Phagocytosis of bacteria as a proposed way to obtain iron is not regulated by iron availability in the parasite; on the contrary, it is decreased. Considering this, it is important to note that the relationship between decreased phagocytosis and decreased hemerythrin expression was previously described [26].

The lack of an inducible iron uptake system has also been described in the parasite *Tritrichomonas foetus* [27]. However, contrary to *N. fowleri*, the obligatory parasite *T. foetus* appears to be able to utilize a wide range of iron sources, including host transferrin or bacterial siderophores. Another potential source of iron for *N. fowleri* can be heme from heme-containing proteins. The ability to utilize exogenous metal-containing porphyrins may be advantageous for protists feeding on bacteria. Moreover, *Naegleria* species phagocyte erythrocytes [28,29], and their ability to degrade hemoglobin using proteases was also described [24]. However, it appears that heme oxygenase is not present in the genome of *N. fowleri*; therefore, it is unlikely to be able to employ this enzyme to obtain iron from hemoglobin, as is the case of *T. foetus* [27]. A potential homologue of bacterial deferrochelatase, a protein able to directly acquire iron from heme [30], was identified as significantly upregulated in iron-deficient conditions by proteomic analysis; however, further investigation is required to clarify the function of this protein. Nevertheless, hemin was shown to partially restore *N. fowleri* growth in very strong iron-deficient conditions. *N. fowleri* requires exogenous porphyrins for growth [31], and the fact that the addition of hemin partly surpasses the conditions of iron starvation can be attributed to a metabolic rearrangement towards heme-dependent pathways.

Proteomic analysis proved to be a valuable resource in determining the cellular changes brought upon *N. fowleri* by iron starvation and gave a foundation for further discoveries shown in this work. The most fundamental finding of this experiment is that mainly cytosolic iron-containing proteins were downregulated. These are components of probably nonessential pathways, such as the phenylalanine degradation pathway, hydrogenase maturation and hydrogen production; the latter is surprisingly shown to take place in the cytosol of *N. gruberi* [32]. On the other hand, mitochondrial iron-dependent proteins were generally unchanged, while some components of the iron-sulfur cluster synthesis machinery were upregulated in iron-deficient conditions, emphasizing the essential role of the iron-dependent respiration process. Consistent with this observation, the mitochondrial iron transporter was slightly upregulated in iron-deficient conditions. Moreover, the carnitine/acylcarnitine carrier was identified in the membrane-enriched proteomic analysis, and its expression was slightly increased in iron-starved cells. This mitochondrial membrane-bound protein is involved in lipid metabolism, which was recently shown to be vital for *N. gruberi* [33]. Another mitochondrial transport protein, the phosphate carrier, was strongly upregulated in iron-deficient conditions, probably as an effort to compensate for impaired respiration and decreased ATP production in the mitochondria of iron-deficient cells.

*N. fowleri*, as well as *N. gruberi,* possesses an AOX in the mitochondria, and our work indicates one of the possible advantages of this respiratory chain element for these organisms. Under iron-deficient conditions, the activity of AOX was significantly increased, even though it required iron, suggesting that this unusual branch of the respiratory chain takes over a portion of the activity in the iron-demanding conventional pathway of respiratory complexes III, IV and cytochrome c, an observation noted in the nonpathogenic amoeba *N. gruberi* previously [34]. This represents a favorable compensation pathway, even though the overall generation of the proton gradient and therefore ATP synthesis is hindered. Considering the reduced efficiency of respiration by iron-starved cells and the presence of lactate dehydrogenase in the genome of *N. fowleri*, it would be reasonable to expect the employment of the lactate dehydrogenase pathway in the regeneration of the cofactor NAD^+^. Such an effect was observed in *Trichomonas vaginalis,* where the cells modulate this pathway as a way of compensating for the metronidazole-induced loss of hydrogenosomal metabolism [35]. However, our metabolomic analysis showed the opposite change; the production of lactate was decreased upon iron starvation, suggesting another, most likely nonessential, function of this pathway that is attenuated due to the hindered rate of overall energy metabolism. Another possible compensatory pathway, ethanol production, is improbable because of the absence of pyruvate decarboxylase or bifunctional aldehyde/alcohol dehydrogenase in the *N. fowleri* genome. The function of cytosolic hydrogenase, which was downregulated in the iron-limited conditions together with its maturation factors, is unknown.

Consistent with our previous study of iron metabolism in *N. gruberi* [34], hemerythrin was dramatically downregulated in iron-deficient conditions, while its expression appeared to be high, based on the intensities obtained in proteomic data from iron-rich cells. The involvement of hemerythrin in the iron metabolism of *Naegleria* is suggestive but unclear. The presence of unbound metals in the cell must be strictly regulated since the imbalance in iron homeostasis can lead to the formation of ROS, mismetallation or other anomalies leading to the incorrect function of proteins. The relationship between hemerythrin and defense against oxidative stress was previously suggested in bacteria [36,37] and so was the role of hemerythrin-related proteins in iron homeostasis [38] or oxygen sensing [39]. One of the basic mechanisms of maintaining the proper intracellular level of metals is the regulation of their acquisition. Since our study shows the lack of such regulation for iron, it is possible that the sequestration of toxic free iron, as well as its storage for use in iron-deficient conditions, is ensured by hemerythrin functioning as a cytosolic iron pool. This hypothesis is supported by the fact that hemerythrin is a nonheme, noniron-sulfur metalloenzyme and is among the most strongly regulated proteins by iron availability, and unlike other proteins, its regulation was detected even at the mRNA level. Another oxygen-binding protein with unclear function, protoglobin, was detected only in iron-sufficient conditions. The role of protoglobin in the metabolism of *N. fowleri* remains to be elucidated.

In conclusion, our work shows that *N. fowleri* possesses only limited capabilities of adaptation to an iron-deficient environment and is surprisingly not able to utilize transferrin as an alternative source of the metal; neither can it induce the rate of iron acquisition under iron starvation, reflecting the lifestyle of a facultative parasite with limited ability of survival in a host. The main strategy of acquiring iron from media appears to be reductive iron uptake. Proteomic analysis of the response to iron starvation demonstrated that a large amount of downregulated proteins in the iron-deficient conditions were nonmitochondrial and nonheme enzymes, except for the heme-containing protein protoglobin, whose expression is regulated at the transcriptional level, unlike the expression of most other affected proteins. Therefore, it can be hypothesized that the fundamental effect of iron deprivation is the degradation of mismetallated cytosolic proteins with a simultaneous increase in iron delivery to mitochondria and induction of iron-sulfur cluster synthesis machinery to ensure essential cell processes. The overall changes in cellular processes in the iron-deficient conditions discussed in this paper are illustrated in Fig 2. These findings are in agreement with our previous study focused on iron metabolism in the nonpathogenic model organism *N. gruberi* [34], where the mitochondrion was shown to be the center of the iron economy. Together, these results show that iron deficiency is a highly unfavorable condition for *N. fowleri*, and targeted interference with its uptake could be an effective method of controlling the propagation or viability of this organism in the host.

**Fig 2.**
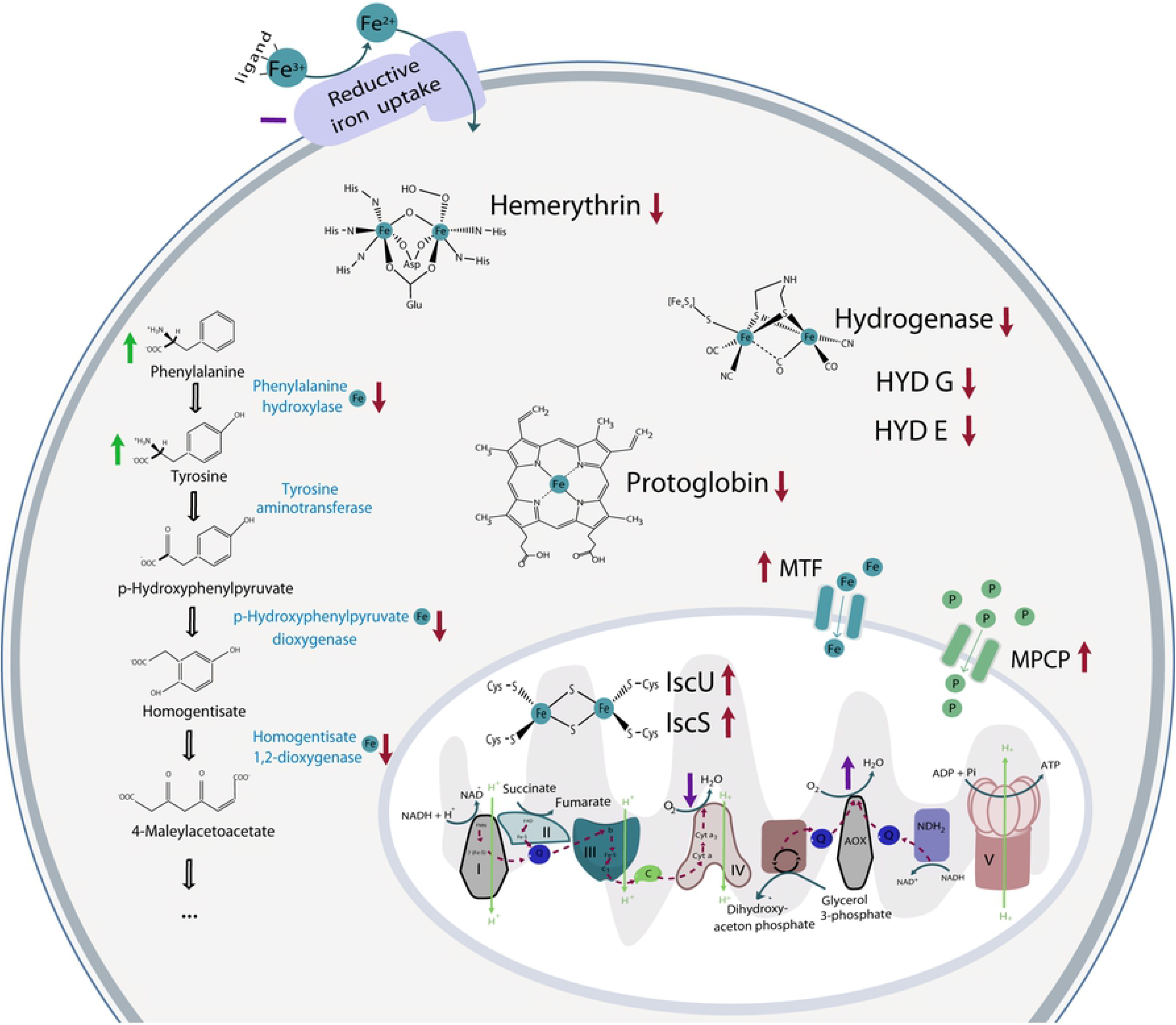
Illustration of the main effects of iron-deficient conditions on the selected cellular processes of *N. fowleri*. The results of proteomic analysis for selected proteins are depicted in red, the results from measured metabolite levels are in green and the assessed enzyme activities are in purple. Upwards pointing arrows stand for increased in the iron-deficient condition, downwards pointing arrows represent decreased in the iron-deficient conditions and dashes mean no significant change in the different iron conditions. Fe represents iron-containing/involving protein/process. C, cytochrome C; MTF, mitoferrin; MPCP, mitochondrial phosphate carrier protein; P, phosphate; Q, ubiquinol/ubiquinone.

## Materials and Methods

### Organisms

*Naegleria fowleri*, strain HB-1, kindly provided by Dr. Hana Pecková (Institute of Parasitology, Biology Center CAS) was maintained in 2% Bacto-Casitone (Difco, USA) supplemented with 10% heat-inactivated fetal bovine serum (Thermo Fisher Scientific, USA), penicillin (100 U/ml) and streptomycin (100 μg/ml) in 25 cm^2^ aerobic cultivation flasks at 37°C. When required, the cells were cultivated for 72 hours with the addition of 25 μM BPS (Sigma-Aldrich, USA), simulating the iron-deficient environment, or with 25 μM Fe-NTA (Sigma-Aldrich, USA) to ensure iron-rich conditions.

### Chelators

The growth dependence of *N. fowleri* on iron availability was assessed using the iron chelators BPS (Sigma-Aldrich, USA), DIP (Sigma-Aldrich, USA) and DFO (Sigma-Aldrich, USA). *N. fowleri* cells were cultivated on 24-well plates in a total volume of 1 ml in a humid chamber. Each chelator and control were tested in biological tetraplicates (final concentrations of 100, 50, 25, 12.5, 6.3, 3.1, 1.6 and 0.8 μM). Cells were cultivated for 48 hours, the plates were placed on ice for 10 min, and the medium was gently pipetted to detach the cells. The number of cells in each sample was counted using a Guava easyCyte 8HT flow cytometer (Merck, Germany). To describe and compare the effect of each chelator, the value of half-maximal inhibitory concentration (IC_50_) was calculated using the online calculator on the AAT Bioquest webpage [40].

### Iron uptake

*N. fowleri* cells grown for 72 hours in iron-rich and iron-deficient conditions were washed with phosphate buffered saline (PBS) (1000 g for 15 min) and transferred to measuring buffer (50 mM glucose; 20 mM HEPES; pH 7.2). Cells were counted using a Guava easyCyte 8HT flow cytometer, and 2.5×10^5^ cells were equally split onto a 24-well plate. To assess iron uptake, cells were supplemented with 2 μM ^55^Fe-citrate, 2 μM ^55^Fe-citrate with 1 mM ascorbate, or 6.3 μM ^55^Fe-transferrin. Samples were incubated at 37°C for 1 hour, and then EDTA was added to a final concentration of 1 mM to chelate extracellular iron. Cells were washed three times by measuring buffer, and the protein concentration was assessed using a BCA kit (Sigma-Aldrich, USA). Samples were diluted to equal concentrations and separated using the Novex Native PAGE Bis–Tris Gel system (4–16%; Invitrogen, USA). The gel was vacuum-dried for 2 hours and autoradiographed using a tritium storage phosphor screen.

### Ferric reductase activity

To assess the activity of *N. fowleri* ferric reductase in different iron environments, a ferrozine assay was used to compare the formation of ferrous iron, as described previously [41]. Cells were grown in iron-rich and iron-deficient environments for 72 hours. Samples containing no cells and samples without Fe-EDTA were used as controls. The cells were washed twice and resuspended in glucose buffer (50 mM glucose, 0.5 mM MgCl_2_, 0.3 mM CaCl_2_, 5.1 μM KH_2_PO_4_, 3 μM Na_2_HPO_4_, pH 7.4). The total amount of proteins was assessed using a BCA kit, and samples were diluted to equal concentrations. All further work was performed with minimum light exposure. Ferrozine was added to a total concentration of 1.3 mM and Fe-EDTA to a concentration of 0.5 mM. Samples were incubated at 37°C for three hours, pelleted (1000 g for 10 min) and the supernatant was used to determine the formation of the colored Fe(II)-ferrozine complex, accompanied by a change in absorbance at 562 nm using 1 cm cuvettes and a UV-2600 UV-VIS spectrophotometer (Shimadzu, Japan).

### Hemin utilization

To assess the ability of *N. fowleri* to utilize hemin, 2×10^4^ cells were cultivated in each well of a 24-well plate in fresh growth medium. Cultures were supplemented with 50 μM BPS, 50 μM hemin, or 50 μM hemin and 50 μM BPS. Cells without chelator or hemin were used as a control. The cell concentration was measured every 24 hours for four days using a Guava easyCyte 8HT flow cytometer as described above.

### Comparative proteomic analysis

*N. fowleri* cells were cultivated in 75 cm^2^ aerobic cultivation flasks in iron-rich and iron-deficient environments. For whole-cell proteomic analysis, cells were washed three times in PBS (1000 g, 15 min, 4°C) and pelleted.

In addition, a membrane-enriched fraction was prepared. Approximately 1.5×10^7^ cells were harvested (1000 g, 15 min, 4°C), washed in 15 ml of sucrose-MOPS buffer (250 mM sucrose, 20 mM 3-morpholinopropanesulfonic acid, pH 7.4, supplemented with complete EDTA-free protease inhibitors, Roche, Switzerland) and resuspended in 4 ml of sucrose-MOPS buffer. The suspension was sonicated using a Q125 sonicator (Qsonica, USA) (45% amplitude, total time of 4 min using 1 s pulse and 1 s pause). The resulting suspension was centrifuged (5000 g, 5 min) to spin down the unlysed cells. The obtained supernatant was further centrifuged (200 000 g, 20 min). The supernatant was discarded, and the membrane-enriched pellet was washed with distilled water.

Label-free proteomic analysis of the samples was assessed using the method described in *Mach et al. 2018* [34], utilizing liquid chromatography coupled with mass spectrometry. The resulting MS/MS spectra were compared with the *Naegleria fowleri* database, downloaded from amoebaDB [42] on 25/7/2017. The set thresholds to filter proteins were Q-value =0, unique peptides detected >2, and the protein had to be identified at least twice in one condition from the six runs. For proteins not identified only in one of the conditions, intensity of 23 was selected as a lowest value to be included, on basis of previous imputation experience. To distinguish significantly downregulated or upregulated proteins in the iron-deficient conditions in S1 and S2 Tables, the threshold of fold change >2.3 and <−2.3 was chosen.

The annotations and identifications of the selected proteins discussed in this work were confirmed using the tool HHPRED [43]. In addition, alignments were constructed using the chosen proteins and their homologues from different organisms: NF0014630 identified as mitochondrial carnitine/acylcarnitine transferase [44], NF0001420 identified as mitochondrial phosphate carrier [45], NF0060430 identified as IscU [46,47] and NF0079420 identified as mitoferrin [48,49]. Alignments with the indicated conserved sequences are shown in S3 Fig and were constructed using Geneious version 11.1.5 software with the muscle alignment tool.

### Comparative transcriptomic analysis

To obtain the transcriptome data of *N. fowleri*, biological pentaplicates of approximately 1×10^6^ cells each were grown in iron-rich and iron-deficient conditions for 72 hours. A High Pure RNA Isolation Kit (Roche, Switzerland) was used to isolate cell RNA, and an Illumina-compatible library was prepared using QuantSeq 3′ mRNA-Seq Library Prep Kit FWD for Illumina (Lexogen, Austria). The RNA concentration was determined using a Quantus fluorometer (Promega, USA), and the quality of RNA was measured on a 2100 Bioanalyzer Instrument (Agilent technologies, USA). Equimolar samples were pooled to 10 pM and sequenced with MiSeq Reagent Kit v3 (Illumina, USA) using 150-cycles on the MiSeq platform. The obtained results were filtered using the P-value of >0.05 followed by analysis on the BlueBee platform with the method DESeq [50].

### Amino acid quantification assay

Approximately 3×10^6^ cells cultivated in iron-rich and iron-deficient environments were harvested by centrifugation (1000 g, 15 min, 4°C), washed with PBS supplemented with cOmplete EDTA-free protease inhibitor, and the total concentration of proteins was measured. Cells were transferred to 1 ml of buffer solution (20 mM Tris, 1 mM MgCl_2_, pH 8, cOmplete EDTA-free protease inhibitor) and sonicated with Sonopuls mini20 (Bandelin, Germany) (90% amplitude, 4°C, total time 120 s, 1 s pulse and 1 s pause). The resulting suspension was mixed at a ratio of 1:4 with ice-cold acetonitrile and maintained overnight at −20°C. Samples were centrifuged (16000 g, 20 min, 4°C) and filtered using Ultrafree Centrifugal Filter Units (Merck Millipore, USA).

Samples were analyzed using liquid chromatography on a Dionex Ultimate 3000 HPLC system with on-line mass spectrometry detection (Thermo Scientific, USA). The separation was achieved using a HILIC column iHILIC-Fusion (150 × 2.1 mm, 1.8 μm particles, 100 Å pore size, HILICON, Sweden). The entire analysis flow rate was 0.3 ml/min, and the column was equilibrated with 100% of solution A (80% acetonitrile in water, 25 mM ammonium formate, pH 4.8) for 3 min. Amino acids were eluted by increasing the gradient of solution B (5% acetonitrile in water, 25 mM ammonium formate, pH 4.8), where 50% of solution B was reached in 7 min. After separation, the column was washed with 80% solution B for 3 min and then equilibrated with 100% solution A for 5 min.

Amino acids were detected by mass spectrometry using the triple quadrupole instrument TSQ Quantiva (Thermo Scientific, USA) in Selected Reaction Monitoring mode. Analytes were ionized using electrospray ionization on an H-ESI ion source and analyzed with positive charge mode with a spray voltage of 3500 V, ion transfer tube temperature of 325°C, and vaporizer temperature of 350°C. All transitions, collision energies, and RF voltages were optimized prior to analysis using appropriate amino acid standards. Each analyte was detected using at least two transitions. Cycle time was set to 1.8 s and both Q1 and Q3 resolutions were set to 0.7 s. To analyze the ion chromatograms and calculate peak areas, Skyline daily version 4.2.1.19004 [51] was used.

### Lactate production

To assess the difference in the intracellular production of lactate, 3×10^6^ cells cultivated in iron-rich or iron-deficient conditions were prepared for analysis in the same way as for quantifying the amino acid content. After incubation with acetonitrile and filtration, the cell sample was dried and resuspended in 100 μl of anhydrous pyridine (Sigma-Aldrich, USA), and 25 μl of a silylation agent (N-tert-butyldimethylsilyl-N-methyl-trifluoroacetamide, Sigma-Aldrich, USA) was added. The sample was incubated at 70°C for 30 min. After incubation, 300 μl of hexane (Sigma-Aldrich, USA) and 10 μl of an internal standard (102 μg/ml 1-bromononane solution in hexane) were added. Selected compounds were analyzed as tert-butyl silyl derivatives.

Samples were analyzed using two-dimensional gas chromatography coupled with mass detection (GCxGC-MS; Pegasus 4D, Leco Corporation, USA) with ChromaTOF 4.5 software. Mass detection was equipped with an EI ion source and TOF analyzer with unite resolution. A combination of Rxi-5Sil (30 m × 0.25 mm, Restek, Australia) and BPX-50 (0.96 m × 0.1 mm, SGE, Australia) columns were used. The input temperature was set to 300°C, the injection volume was 1 μl in spitless mode, and constant helium flow of 1 ml/min, modulation time 3 s (hot pulse 1 s) and modulation temperature offset with respect to the secondary oven 15°C were used. The temperature program applied on the primary oven was 50°C (hold 1 min), which was increased by the rate of 10°C/min to a final temperature of 320°C (hold 3 min). The temperature offset applied on the secondary column was +5°C.

### Cell respiration

Five biological replicates of 3×10^6^ *N. fowleri* cells grown for 72 hours in iron-rich and iron-deficient conditions were washed twice and resuspended in 1 ml of glucose buffer, and the protein concentration was assessed using a BCA kit. Total cell respiration was measured as the decrease in oxygen concentration using an Oxygen meter model 782 (Strathkelvin instruments, UK) with Mitocell Mt 200 cuvette of total volume of 700 μl at 37°C. Measurements were carried out with 5×10^5^ cells for 5 min, after which potassium cyanide was added to a final concentration of 4 mM to block complex IV, and after 5 min, salicyl hydroxamic acid was added to a final concentration of 0.2 mM to completely block AOX. Values gained after the addition of potassium cyanide and salicyl hydroxamic acid were subtracted to acquire canonical respiratory chain and AOX activity, respectively.

### *N. fowleri* bacterial phagocytosis

Approximately 3×10^6^ cells cultivated in iron-rich and iron-deficient conditions were washed in cultivation flasks by replacing the growth medium with 10 ml of PBS warmed to 37°C and resuspending in 7 ml of 37°C PBS. To assess the ability of the cells to phagocytose, 150 μl of pHrodo™ Green *E. coli* BioParticles™ Conjugate for Phagocytosis (Thermo Fisher Scientific, USA) was added, and the cells were incubated for 3 hours at 37°C. Following incubation, the cells were washed with PBS, detached on ice for 15 min, and the fluorescence caused by the phagocytosed particles was consecutively analyzed using a Guava easyCyte 8HT flow cytometer as described above using a 488 nm laser and a Green-B 525/30 nm detector. A negative control (without the addition of BioParticles) was used to determine an appropriate threshold for *N. fowleri* cells. BioParticles resuspended in PBS were measured in the same way to determine the background noise and gave a negligible signal. The effect of different iron availability on the efficiency of phagocytosis of *N. fowleri* was established as the percentage of cells in culture that had increased fluorescence.

To visualize the ability of *N. fowleri* to phagocytize, live amoebae incubated with fluorescent *E. coli* were imaged with a Leica TCS SP8 WLL SMD-FLIM microscope (Leica, Germany) equipped with an HC PL APO CS2 63x/1.20 water objective with 509 nm excitation and 526 nm-655 nm for HyD SMD and a PMT detector for brightfield imaging. Images were processed using LAS X 3.5.1.18803 (Leica, Germany).

## Acknowledgements

Special thanks to Ivánek for fruitful biochemical discussions.

## Supporting information

**S1 Fig. (A). Lack of ^55^Fe-transferrin uptake in *N. fowleri*, cultivated in iron-rich and iron-deficient conditions.** The uptake of transferrin-bound iron was assessed by incubation of *N. fowleri* with ^55^Fe-transferrin. Tf, pure ^55^Fe-transferrin; Fe *N. fowleri* cultivated in iron-rich conditions for 72 hours, consecutively incubated with ^55^Fe-transferrin for 1 hour; BPS, *N. fowleri* cultivated in iron-deficient conditions for 72 hours, consecutively incubated with ^55^Fe-transferrin for 1 hour; Ctr, iron uptake control of *N. fowleri* culture cultivated in iron-deficiency incubated with ^55^Fe(III)-citrate for 1 hour. The utilization of iron was analyzed by blue native electrophoresis as described in the Methods section. **(B) Mechanism of ferric iron uptake involves the reductive step.** *N. fowleri* culture was incubated for 1 hour with ^55^Fe(III)-citrate with and without the addition of 0.2 mM BPS. Incorporation of ^55^Fe(III)-citrate to cellular proteins was higher in the sample without the presence of BPS, indicating that a reductive iron uptake mechanism takes place. Several distinct signals on the lower part of +BPS probably correspond to residues of BPS complexed with ferrous iron radionuclides. The utilization of iron was analyzed by blue native electrophoresis as described in the Methods section. −BPS, cell sample without addition of BPS chelator; +BPS, cell sample with the addition of BPS chelator. **(C) Loading control of *N. fowleri* cell lysate for Western blot analysis of hemerythrin expression.** Ponceau S loading stain shows equal protein concentrations of loaded samples of *N. fowleri* cultivated in iron-deficient (BPS) and iron-rich (Fe) conditions.

**S2 Fig. *N. fowleri* cell phagocytosis.** *N. fowleri* cells phagocytizing several fluorescently modified pHrodo™ Green *E. coli* BioParticles™ (blue). Images were acquired with a Leica TCS SP8 WLL SMD-FLIM microscope (Leica, Germany) equipped with an HC PL APO CS2 63x/1.20 water objective with 509 nm excitation and 526 nm-655 nm for HyD SMD and a PMT detector for brightfield imaging. Images were processed using LAS X 3.5.1.18803 (Leica, Germany).

**S3 Fig. Alignments of *N. fowleri* proteins with homologues from other organisms** (A) Alignment of *N. fowleri* NF0060430 with the IscU proteins from *Homo sapiens*, *Saccharomyces cerevisiae* and *T. brucei*. Red arrows point to the conserved cysteine required for iron-sulfur cluster assembly, based on a previous study [46]. The red rectangle denotes the conserved LPPVK motif of the IscU proteins [47]. (B) Alignment of *N. fowleri* NF0079420 with the mitoferrin proteins of *Trypanosoma brucei*, *Leishmania mexicana*, *Saccharomyces cerevisiae* and *Homo sapiens*. Red arrows point to the sequence motif Px(D/E)xx(K/R)x(K/R), and yellow circles mark residues in contact with substrate, according to a previous study [48]. Conserved histidine residues responsible for iron transport are marked with blue stars [49]. (C) Alignment of *N. fowleri* NF0001420 with the mitochondrial phosphate carriers of *Saccharomyces cerevisiae*, *Homo sapiens* and *Arabidopsis thaliana*. Red arrows point to residues important for the phosphate transport activity, according to a previous study [45]. (D) Alignment of *N. fowleri* NF0014630 with mitochondrial carnitine/acylcarnitine transferases of *Saccharomyces cerevisiae*, *Arabidopsis thaliana*, *Homo sapiens* and *Caenorhabditis elegans*. The red rectangle denotes the signature motifs Px(D/E)xx(R/K)x(R/K), and the arrows point to conserved residues, according to a previous study [44].

**S1 Table. Comparison of whole-cell proteomes of *N. fowleri* in iron-rich and iron-deficient environments.** List of whole-cell proteomes of *N. fowleri* compared in iron-rich and iron-deficient conditions sorted into four sheets: raw data, all detected proteins, significantly upregulated proteins and significantly downregulated proteins in iron-deficient conditions. Proteins with >2.3 (denoting upregulated in iron-deficient conditions) or <−2.3-fold change (denoting downregulated in iron-deficient conditions) are regarded as significantly regulated. Proteins were annotated from amoebaDB [42] on 25/7/2017, and manual annotation was performed for selected proteins, as described in the Methods section. Probable iron-containing proteins of the significantly downregulated and upregulated proteins are highlighted in yellow.

**S2 Table. Comparison of membrane proteomes of *N. fowleri* in iron-rich and iron-deficient environments.** List of membrane-enriched proteomes of *N. fowleri* compared in iron-rich and iron-deficient conditions sorted into four sheets: raw data, all detected proteins, significantly upregulated proteins and significantly downregulated proteins in iron-deficient conditions. Proteins with >2.3 (denoting upregulated in iron-deficient conditions) or <−2.3-fold change (denoting downregulated in iron-deficient conditions) are regarded as significantly regulated. Proteins were annotated from amoebaDB [42] on 25/7/2017, and manual annotation was performed for selected proteins, as described in the Methods section.

**S3 Table. Comparison of transcriptomes of *N. fowleri* in iron-rich and iron-deficient environments.** List of genes significantly downregulated and upregulated in iron-deficient conditions. Raw data are included, and manual annotation was performed for selected proteins, as described in the Methods section.

